# Control of neuronal terminal differentiation through cell context-dependent CFI-1/ARID3 functions

**DOI:** 10.1101/2022.07.04.498728

**Authors:** Yinan Li, Jayson J. Smith, Filipe Marques, Anthony Osuma, Hsin-Chiao Huang, Paschalis Kratsios

## Abstract

ARID3 transcription factors are expressed in the nervous system, but their functions and mechanisms of action are largely unknown. Here, we generated *in vivo* a genome-wide binding map for CFI-1, the sole *C. elegans* ARID3 ortholog. We identified 6,396 protein-coding genes as putative direct targets of CFI-1, most of which (77%) are expressed in post-mitotic neurons and encode terminal differentiation markers (e.g., neurotransmitter receptors, ion channels, neuropeptides). To gain mechanistic insights, we focused on two neuron types. In sensory neurons (IL2 class), CFI-1 exerts a dual role: it acts directly to activate, and indirectly to repress, distinct terminal differentiation genes. In motor neurons, however, CFI-1 acts directly as a repressor, continuously antagonizing three transcriptional activators (UNC-3/Ebf, LIN-39/Hox4-5, MAB-5/Hox6-8). By focusing on a glutamate receptor gene (*glr-4/GRIK1*), we found CFI-1 exerts its repressive activity through proximal binding to the *glr-4* locus. Further, the core DNA binding domain of CFI-1 is partially required for *glr-4* repression in motor neurons. Altogether, this study uncovers cell context-dependent mechanisms through which a single ARID3 protein controls the terminal differentiation of distinct neuron types.

## INTRODUCTION

Members of the ARID family of proteins are found in plants, yeast, fungi, and invertebrate and vertebrate animals (Kortschak et al., 2000, Patsialou et al., 2005, Wilsker et al., 2002, Wilsker et al., 2005). ARID family proteins are expressed either ubiquitously or in a tissue-specific fashion and control various biological processes, such as cell proliferation, differentiation, and embryonic patterning (Wilsker et al., 2002, Wilsker et al., 2005). Additionally, mutations in ARID family proteins are associated with cancer and several neurodevelopmental disorders (Shang et al., 2015, Bramswig et al., 2017, Kosho et al., 2014, Miyake et al., 2014, Smith et al., 2016, Lin et al., 2014).

Humans possess fifteen ARID family proteins, divided into seven subfamilies (ARID1-5, JARID1-2) based on the degree of sequence similarity (Wilsker et al., 2005). The **A**T-**R**ich **I**nteraction **D**omain (ARID), after which the family is named, was first identified in ARID3 proteins. These bind DNA in a sequence-specific manner, prefer AT-rich sequences, and are known to function as transcription factors (Wilsker et al., 2005). The ARID5 subfamily also encodes transcription factors (Patsialou et al., 2005, Wilsker et al., 2002), but the remaining five subfamilies (ARID1-2, ARID4, JARID1-2) encode proteins that bind DNA in a non-sequence-specific manner (Patsialou et al., 2005). For example, ARID1A, ARID1B, and ARID2 constitute subunits of the SWI/SNF (BAF/PBAF) chromatin-remodeling complex that can move and/or eject nucleosomes. Although the precise functions of all ARID proteins are not known, accumulating evidence suggests they can act either as positive or negative regulators of gene transcription, or as components of chromatin-remodeling complexes (Kortschak et al., 2000, Patsialou et al., 2005, Wilsker et al., 2002, Wilsker et al., 2005).

Mouse *Bright* (*Arid3a*) and *Drosophila dead ringer* (*retained*) are the founding members of the ARID family, and belong to the ARID3 subfamily, which is specific to metazoans. Single orthologs exist in *C. elegans* (CFI-1) and *Drosophila (dead ringer),* whereas mammals contain three orthologs (*ARID3A-C)* (Fig. 1A) (Kortschak et al., 2000, Wilsker et al., 2005, Gregory et al., 1996, Shaham and Bargmann, 2002). A defining feature of ARID3 proteins is the <u>e</u>xtended ARID (eARID) domain, a ∼40 residue-long region next to the core ARID domain (Wilsker et al., 2002, Wilsker et al., 2005) (Kortschak et al., 2000). Structural studies showed both ARID and eARID domains contact DNA (Iwahara and Clubb, 1999, Iwahara et al., 2002).

**Figure 1.**
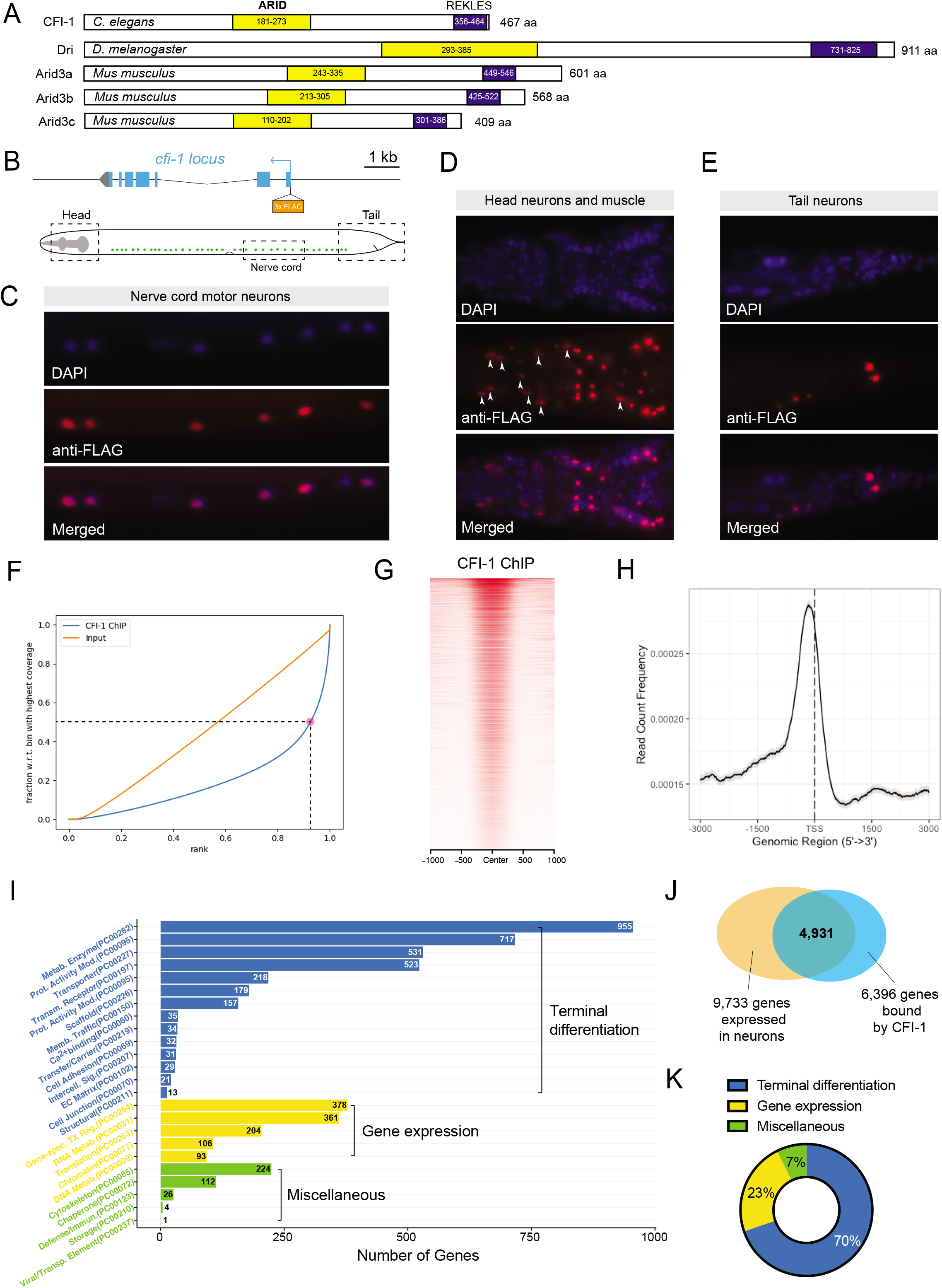
Mapping genome-wide CFI-1 binding with ChIP-seq. **A:** Schematics of the coding sequences of CFI-1 and its *Drosophila* homolog *Dead ringer (Dri)* and Mouse homologs Arid3a-c. The ARID (yellow) and REKLES (blue) domains are highlighted. **B:** (Doitsidou et al.) Diagram of the *3xflag::cfi-1* allele. The endogenous CFI-1 proteins are tagged with 3xFLAG which are inserted immediately after the ATG. (Bottom) Schematic of *C. elegans* with dashed line boxes highlighting regions where immunostaining images are shown in C-E. **C-E:** The expression of 3xFLAG::CFI-1 fusion protein is confirmed by immunostaining (DAPI, blue; anti-FLAG, red) in the ventral cord motor neurons (C), muscles (indicated by arrowheads) and neurons in the head (D), and neurons in the tail (E). **F:** Fingerprint plot indicating localized, strong enrichment of CFI-1 binding events in the genome. Specifically, when counting the reads contained in 86% of all genomic bins, only 50% of the maximum number of reads are reached, which indicates 14% of the genome contain half of total reads. **G:** Heatmap of CFI-1 ChIP-seq signal around 1.0 kb of the center of the binding peak. **H:** Summary plot of CFI-1 ChIP-seq signal with a 95% confidence interval (grey area) around 3.0 kb of the transcription start site (Kadkhodaei et al.). The average signal peak is detected at ∼140 bp upstream of the TSS. **I:** Graph summarizing protein class ontology analysis of global CFI-1 target genes identified by ChIP-seq. A total number of 7,995 genes are analyzed, 4,984 of which hit known protein class terms. **J:** Pie chart summarizing three main categories of genes bound by CFI-1. **K**: Venn diagram showing that 77.1% of the protein-coding genes (4,931 out of 6,396) that are bound by CFI-1 are expressed in the nervous system based on RNA-Seq data from the CenGEN project.

ARID3 proteins have several early developmental roles, as identified by genetic studies. Mice lacking *Bright/Arid3a* display early embryonic lethality due to defects in hematopoiesis (Webb et al., 2011). *Bright/Arid3a* is best studied in B cell lineages, where it acts as an activator and increases immunoglobulin transcription(Herrscher et al., 1995, Ratliff et al., 2014, Webb et al., 2011, Webb et al., 1998). However, *Bright/Arid3a* is also critical for embryonic stem cell differentiation (An et al., 2010, Popowski et al., 2014, Rhee et al., 2014). In this context, it can act either as an activator or repressor of gene expression (Rhee et al., 2014). Similar to mice lacking *Bright/Arid3a,* null mutants for *dead ringer* in *Drosophila* display early lethality (Shandala et al., 1999, Shandala et al., 2002)*. Dead ringer* is essential for anterior-posterior patterning and muscle development in the fly embryo, and can act either as an activator or repressor of gene transcription(Hader et al., 2000, Shandala et al., 1999, Valentine et al., 1998). Lastly, *Arid3a* and *Arid3b* have been associated with tumorigenesis by acting as direct inducers of cell cycle regulators (Lestari et al., 2012, Saadat et al., 2021).

Genetic studies in *Drosophila* and *C. elegans* have also identified late developmental roles of ARID3 proteins in the nervous system. In *Drosophila* larvae, *dead ringer* is expressed in distinct neuron types and controls axonal pathfinding (Ditch et al., 2005, Shandala et al., 2003, Sibbons, 2004), though its downstream targets in neurons -and thus, whether *dead ringer* behaves as an activator or repressor – remain unknown. In *C. elegans*, *cfi-1* is selectively expressed in head muscle and several neuron types: the IL2 sensory neurons, the AVD, PVC, and LUA interneurons, and various classes of motor neurons (Shaham and Bargmann, 2002, Li et al., 2020). Because *C. elegans* animals lacking *cfi-1* are viable (Shaham and Bargmann, 2002), these mutant strains provided a glance into the potential functions of ARID3 proteins during post-embryonic life. Candidate approaches that examined a handful of effector genes encoding neurotransmitter (NT) biosynthesis proteins and receptors suggested that CFI-1 acts as an activator of gene expression both in sensory neurons (IL2) and interneurons (AVD, PVC) (Shaham and Bargmann, 2002, Zhang et al., 2014, Ahn et al., 2022). However, in ventral nerve cord motor neurons, CFI-1 is thought to act as a repressor of the glutamate receptor gene *glr-4/GRIK1* (Kerk et al., 2017). The molecular mechanisms underlying the differential activities of CFI-1 (activator versus repressor) in these distinct neuron types remain unknown. Elucidating such mechanisms in *C. elegans* may provide clues as to how CFI-1 orthologs in other species control cell differentiation. Lastly, unbiased approaches to identify the *in vivo* targets of CFI-1 (and any other ARID3 protein) in the nervous system are currently lacking, preventing a comprehensive understanding of the neuronal functions controlled by ARID3 transcription factors.

Here, we performed chromatin immunoprecipitation for CFI-1 followed by sequencing (ChIP-Seq). By generating an *in vivo* binding map on the *C. elegans* genome, we identified 6,396 protein-coding genes as putative direct targets of CFI-1, the majority of which (77%) are expressed in post-mitotic neurons. Gene ontology analysis suggests CFI-1 is primarily involved in the process of neuronal terminal differentiation. To gain mechanistic insight into how CFI-1 controls the terminal differentiation of different neuron types, we focused on head sensory neurons (IL2 class) and nerve cord motor neurons (DA, DB, VA, and VB classes). In sensory IL2 neurons, CFI-1 exerts a dual role: it acts directly to activate and indirectly to repress distinct terminal differentiation genes (e.g., NT receptors, ion channels). In nerve cord motor neurons, however, CFI-1 acts directly to repress expression of the glutamate receptor gene *glr-4/GRIK1*. CRISPR/Cas9-mediated mutagenesis of endogenous CFI-1 binding sites suggests proximal binding to the *glr-4* locus is necessary for repression, advancing our understanding of ARID3-mediated gene repression. Importantly, the core DNA binding domain of CFI-1 is partially required for *glr-4* repression in motor neurons. Altogether, this study offers mechanistic insights into cell context-dependent functions of CFI-1 (ARID3), a critical regulator of the terminal differentiation program of distinct neuron types.

## RESULTS

### A map of CFI-1/ARID3 binding on the *C. elegans* genome

To identify CFI-1 binding events, we first generated an endogenous reporter strain through in-frame insertion of the *flag* epitope sequence (*3xflag*) immediately after the *cfi-1* start codon (Fig. 1B). Immunostaining against FLAG on adult *3xflag::cfi-1* animals showed nuclear expression in head muscle cells, as well as in neurons of the head, ventral nerve cord, and tail regions (Fig. 1C-E), indicating this reporter allele faithfully recapitulates the known expression pattern of *cfi-1* (Kerk et al., 2017, Shaham and Bargmann, 2002). Unlike *cfi-1* null animals (Shaham and Bargmann, 2002), homozygous *3xflag::cfi-1* animals do not display any defects in posterior touch response (Supplementary Fig. 1). This suggests that insertion of the *3xflag* sequence does not alter *cfi-1* gene function. We therefore conducted ChIP-Seq using a FLAG antibody on homozygous *3xflag::cfi-1* animals at the third larval stage (L3), as all *cfi-1-*expressing cells are generated by this stage.

Our ChIP-Seq experiment revealed strong enrichment of CFI-1 binding in the *C. elegans* genome, identifying 14,806 unique binding peaks (q-value cutoff: 0.05) (Fig. 1F-G). The CFI-1 peaks are predominantly located between 0 and 3kb upstream of transcription start sites (Fig. 1H), suggesting CFI-1 acts at promoter and enhancer regions to regulate gene expression. Altogether, ChIP-Seq for CFI-1 generated the first *in vivo* binding map of an endogenously tagged ARID3 protein, offering an opportunity to comprehensively identify the biological processes controlled by CFI-1.

### The majority of CFI-1/ARID3 target genes encode neuronal terminal differentiation markers

Subsequent bioinformatic analysis of the 14,806 CFI-1 binding peaks revealed 6,396 protein-coding genes as putative CFI-1 targets (see Materials and Methods). Because the majority of *cfi-1-*expressing cells are neurons (Fig. 1C-E) (Kerk et al., 2017, Shaham and Bargmann, 2002), we reasoned that a significant portion of the 6,396 protein-coding genes may be expressed in the nervous system. To test this, we used available single-cell expression profiles (CeNGEN project: www.cengen.org) for all known *cfi-1*-expressing neurons (IL2, URA, AVD, PVC, LUA, DA, DB, VA, VB, DD, VD), and indeed found that 77.1% of the global CFI-1 targets (4,931 out of 6,396) are expressed in these neurons (Fig. 1J**, Supplementary file 1**). To gain insights into the biological functions of CFI-1, we conducted gene ontology (GO) analysis with PANTHER (Mi et al., 2013). Strikingly, the majority of CFI-1 target genes (∼70%) encodes proteins essential for neuronal terminal differentiation (e.g., NT receptors, transporters, ion channels, transmembrane receptors, cell adhesion molecules) (Fig. 1I-K**, Supplementary file 2**). The second largest category (23% of CFI-1 targets) contains transcription factors, chromatin factors, as well as proteins involved in DNA/RNA metabolism (Fig. 1I-K), suggesting CFI-1 can affect gene expression indirectly through these factors. Altogether, the downstream targets identified via our unbiased approach suggest that CFI-1 plays a prominent role in neuronal terminal differentiation.

### CFI-1/ARID3 acts directly to activate terminal differentiation genes in IL2 sensory neurons

Although *cfi-1* is expressed in several neuron types, a handful of CFI-1 target genes have only been identified in head sensory neurons of the IL2 class (Fig. 2A-B) (Shaham and Bargmann, 2002, Zhang et al., 2014). In these neurons, genetic experiments suggested CFI-1 influences gene expression both positively and negatively (Shaham and Bargmann, 2002, Zhang et al., 2014). CFI-1 activates various terminal differentiation genes (e.g., *cho-1/ChT, unc-17/VAChT, gcy-19* [receptor-type guanylate cyclase]*, klp-6* [kinesin-like protein], *unc-5* [netrin receptor]), and represses expression of two ion channel-encoding genes (*pkd-2/Polycystin-2 like 1* [PKD2L1] and *lov-1/Polycystin-1 like 3* [PKD1L3]) associated with polycystic kidney disease (Zheng et al., 2018b). It remained unknown, however, whether CFI-1 acts directly or indirectly to control these genes. Leveraging our ChIP-Seq dataset, we found that CFI-1 binds directly to all known terminal differentiation genes (e.g., *cho-1/ChT, unc-17/VAChT)* that require *cfi-1* gene activity for their activation in IL2 neurons (Fig. 2C). However, we did not detect any binding in the *cis*-regulatory regions of genes repressed by *cfi-1* (*pkd-2, lov-1*) (Fig. 2D). Altogether, biochemical evidence (ChIP-Seq) combined with genetic studies (Shaham and Bargmann, 2002, Zhang et al., 2014) strongly suggest that in IL2 sensory neurons, CFI-1 exerts a dual role: it directly activates a set of terminal differentiation genes, but indirectly (via intermediary factors) represses the expression of ion channel-encoding genes (*pkd-2/Polycystin-2* and *lov-1/Polycystin-1 like)* (Fig. 2B).

**Figure 2.**
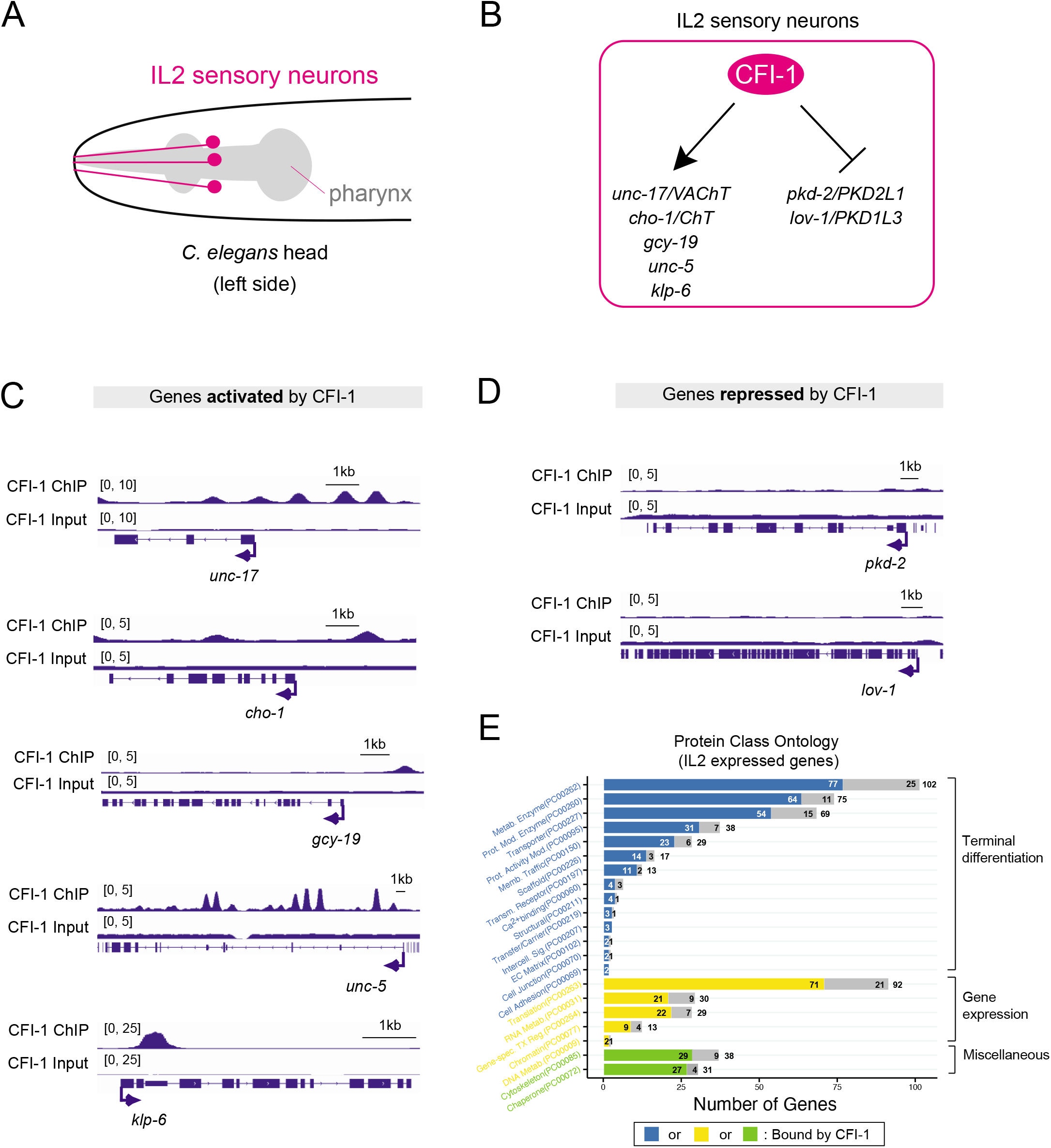
CFI-1 directly activates terminal differentiation genes in the IL2 sensory neurons. **A:** Diagram showing the location of the IL2 sensory neurons. **B:** Model summarizing known targets of CFI-1 in the IL2 sensory neurons. **C:** CFI-1 binding signals at the loci of genes activated by CFI-1 in the IL2 neurons. **D:** CFI-1 binding signals at the loci of genes repressed by CFI-1 in the IL2 neurons. **E:** Protein class ontology analysis of genes expressed in IL2 sensory neurons binned by CFI-1 binding based on ChIP-seq data. Bar colors correspond to colors in Fig 1I: Blue, genes involved in terminal differentiation; yellow, genes involved in regulating gene expression; green, miscellaneous genes.

We next undertook an unbiased approach to investigate whether genes expressed in mature IL2 neurons are also bound by CFI-1. To this end, we used available single-cell expression profiles (CeNGEN project: www.cengen.org) and selected the most highly expressed genes (top 1,000) in IL2 neurons. Strikingly, most of these genes (∼70%) are bound by CFI-1 (Fig. 2E). Among them, GO analysis revealed an overrepresentation of terminal differentiation genes (Fig. 2E**, Supplementary file 3**). This analysis provides biochemical evidence to support the idea that CFI-1 directly activates scores of terminal differentiation genes in IL2 neurons.

### CFI-1/ARID3 acts directly to repress endogenous *glr-4/GRIK1* expression in cholinergic motor neurons

Our findings in IL2 sensory neurons suggest CFI-1 is a direct activator and indirect repressor of gene expression (Fig. 2B-D). Next, we interrogated the function of CFI-1 in cholinergic nerve cord motor neurons that control locomotion (Fig. 3A). Using an endogenous reporter allele (Li et al., 2020), we found that *cfi-1* is selectively expressed in 29 cholinergic motor neurons (of the DA, DB, VA, VB classes) located in the mid-body region (Fig. 3A, Supplementary Fig. 2). Next, we asked whether in these neurons CFI-1 functions as an activator of gene expression. To this end, we examined for *cfi-1* dependency five available motor neuron-specific terminal differentiation markers (*twk-40, twk-43* [TWiK potassium channels]*; ncs-2* [neuronal calcium sensor]*; npr-29* [neuropeptide]*; dbl-1* [Bmp-like]) (Li and Kratsios, 2021). These showed no difference in expression in motor neurons of *cfi-1(-)* mutants (Supplementary Fig. 3). Consistently, a previous study also found that three other terminal differentiation markers (*acr-5* [acetylcholine receptor], *del-1* [SCNN1 sodium channel], *inx-12* [gap junction protein]) are not affected in motor neurons of *cfi-1(-)* mutants (Kerk et al., 2017). Interestingly, our ChIP-seq data indicate that these eight genes are all bound by CFI-1 (Supplementary Fig. 3), raising the possibility of CFI-1 operating redundantly with other transcription factors to activate expression of terminal differentiation genes in motor neurons.

**Figure 3.**
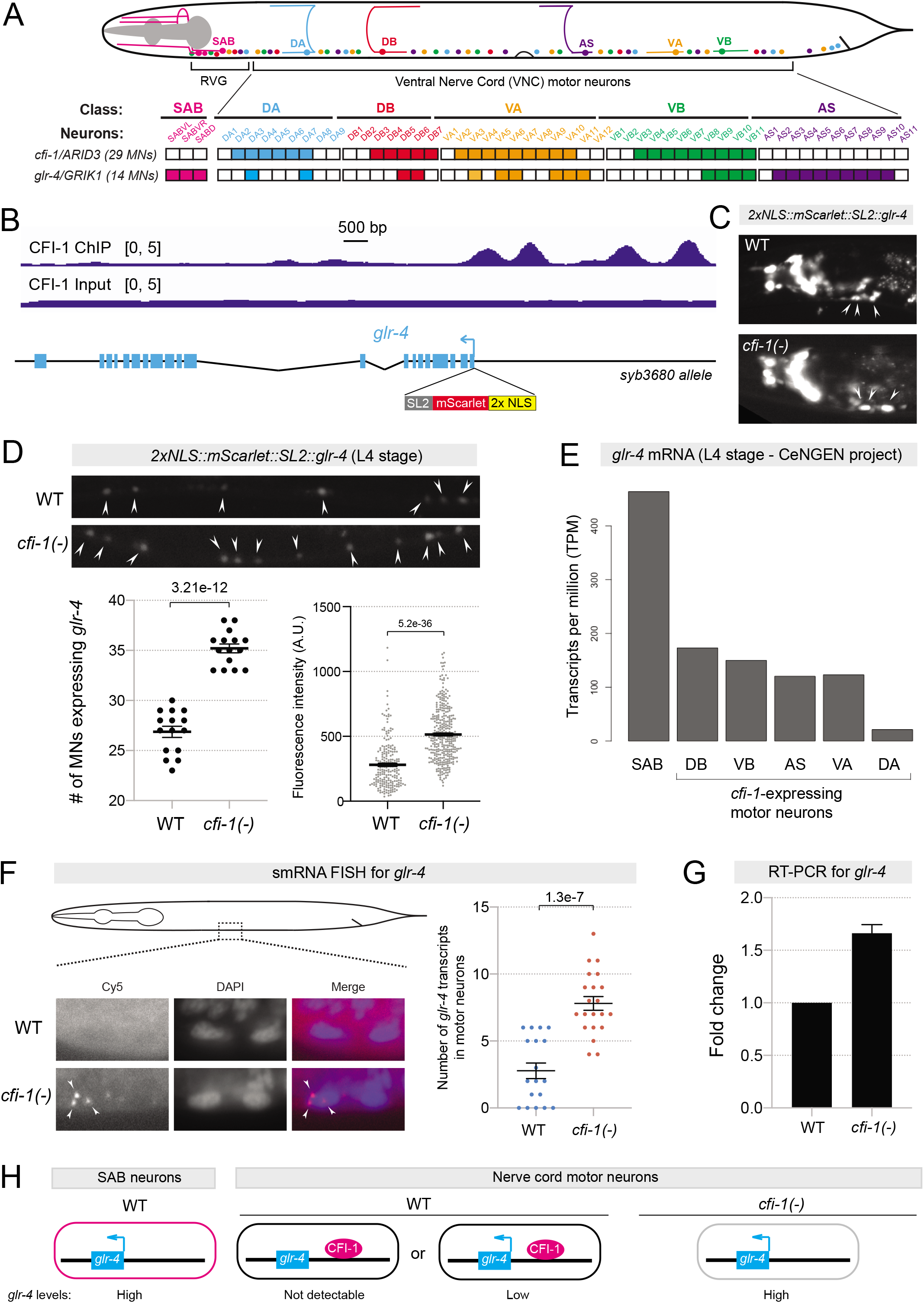
CFI-1/Arid3a represses *glr-4*/GRIK1 in the ventral cord motor neurons. **A:** Summary of the endogenous expression patterns of cfi-1 and glr-4 in the five cholinergic nerve cord motor neuron subtypes and the SAB neurons in the retrovesicular ganglion (RVG). The expression patterns are determined by colocalization with neuron subtype-specific reporters. Filled boxes represent positive expression, while empty boxes indicate no detectable expression. **B:** CFI-1 binding signal at the *glr-4* locus. (Bottom) Design of an endogenously tagged, nuclear localized fluorescent reporter allele of *glr-4* (*2xNLS::mScarlet::SL2::glr-4*). The reporter cassette is inserted immediately downstream of the ATG of *glr-4* with CRISPR/Cas9. **C:** Representative fluorescent micrographs showing that the expression of 2xNLS::mScarlet::SL2::GLR-4 is unaffected in SAB neurons (arrowheads) in *cfi-1(-)* mutants. Bright white signal to the left of the SAB neurons indicates *glr-4* expression in head neurons. **D:** *2xNLS::mScarlet::SL2::glr-4* expression in the ventral cord motor neurons (indicated by arrowheads in the representative images) in WT and *cfi-1(-)* animals at L4 stage. Ectopic expression of *glr-4* was detected in *cfi-1(-)* mutants. (Bottom) Each dot in the quantification graph of total number of motor neurons (left) represents an individual animal. Each dot in the fluorescence intensity quantification graph (right) represents an individual motor neuron that shows *glr-4* expression. For simplicity, only motor neurons in the anterior VNC are included in the fluorescence intensity quantification. p-values are indicated in the graphs. N ≥ 15. **E:** RNA-seq data from the CeNGEN project showing expression of *glr-4* transcripts in SAB, DB, VB, AS, VA, and DA motor neurons. **F:** Single molecule fluorescent *in situ* hybridization (smFISH) shows mild ectopic expression of *glr-4* transcripts in the motor neurons of L1 *cfi-1(-)* animals. Left: represented images showing ectopic *glr-4* mRNA molecules (Cy5, red) in the nucleus (DAPI, blue) of a motor neuron in *cfi-1(-)* mutants. Right: quantification of the number of *glr-4* transcripts detected in the anterior ventral nerve cord in WT and *cfi-1(-)* animals. p-value is indicated in the graph. N ≥ 18. **G:** RT-PCR from whole worm lysates showing upregulation of *glr-4* transcripts in *cfi-1(-)* mutants. **H:** Models summarizing the repressive regulation of glr-4 expression by CFI-1 in the SAB neurons versus ventral nerve cord motor neurons.

The terminal differentiation gene *glr-4* (ortholog of human GRIK1 [glutamate inotropic receptor kainite type subunit 1]) is the only known CFI-1 target in motor neurons, where it is negatively regulated by CFI-1 (Kerk et al., 2017), providing an opportunity to obtain mechanistic insights into how ARID3 proteins mediate gene repression in the nervous system. We therefore carried out an in-depth investigation focused on *glr-4*, as detailed below.

Because previous studies employed transgenic reporters (Brockie et al., 2001, Kerk et al., 2017), the endogenous expression pattern of *glr-4* in *C. elegans* neurons remained unclear. We therefore generated an endogenous *glr-4* reporter allele by inserting the *2xNLS::mScarlet::SL2* cassette immediately after the start codon (see Materials and Methods) and established the *glr-4* expression pattern in motor neurons with single-cell resolution (Fig. 3B). Consistent with previous studies (Brockie et al., 2001, Kerk et al., 2017), we observed high levels of *glr-4* (*mScarlet*) expression in head neurons, as well as in SAB motor neurons that innervate head muscle (Fig. 3A, C). This endogenous reporter also revealed new sites of expression. In the nerve cord of WT animals, we detected low levels of *glr-4* (*mScarlet*) expression in 14 of the 29 *cfi-1*-expressing motor neurons, as well as in AS motor neurons, which do not express *cfi-1* (Fig. 3A-B,D, Supplementary Fig. 2). Further, the observed levels of *glr-4* expression in head (SAB) and ventral cord motor neurons were independently confirmed by available scRNA-Seq data (CeNGEN project: www.cengen.org) (Fig. 3E).

Next, we tested whether endogenous *glr-4* expression in motor neurons depends on *cfi-1* gene activity. Indeed, expression of the *glr-4* (*mScarlet*) reporter allele is increased in motor neurons of *cfi-1* loss-of-function mutants, as we observed a higher number of cells expressing *glr-4,* and at higher levels, compared to WT motor neurons (Fig. 3D). No effects on *glr-4* were observed in SAB neurons, as they do not express *cfi-1* (arrowheads in Fig. 3C). Moreover, we performed single-molecule mRNA fluorescent *in situ* hybridization (sm mRNA FISH) in WT and *cfi-1* mutant animals. Loss of *cfi-1* led to increased levels of *glr-4* mRNA in nerve cord motor neurons (Fig. 3F). Lastly, these results were corroborated by RT-PCR in WT and *cfi-1* mutant animals (Fig. 3G). Altogether, we conclude CFI-1 limits the endogenous expression of *glr-4/GRIK1* in nerve cord motor neurons. In WT animals, we can either detect low or no *glr-4* expression in motor neurons, whereas loss of *cfi-1* results in robust *glr-4* expression in these cells (Fig. 3H). Because ChIP-Seq revealed extensive CFI-1 binding immediately upstream of the *glr-4* locus (Fig. 3B), we propose CFI-1 acts as a direct repressor of *glr-4/GRIK1*.

### *cfi-1/ARID3* is required to maintain *glr-4* repression in nerve cord motor neurons

The *cfi-1*-expressing motor neurons (DA, DB, VA, VB) are generated in two waves (DA/DB are born during embryogenesis; VA and VB at larval stage 1 [L1]) (Fig. 4A-B). Although all *cfi-1*-expressing motor neurons have been generated by L2, we found no *glr-4* (*mScarlet)* expression in WT animals at this stage. However, we observed a progressive increase in the number of WT motor neurons expressing low levels of *glr-4* (*mScarlet)* at subsequent stages (L3, L4, Day 2 [D2] adult), indicating a correlation between *glr-4* expression and motor neuron maturation (Fig. 4A).

**Figure 4.**
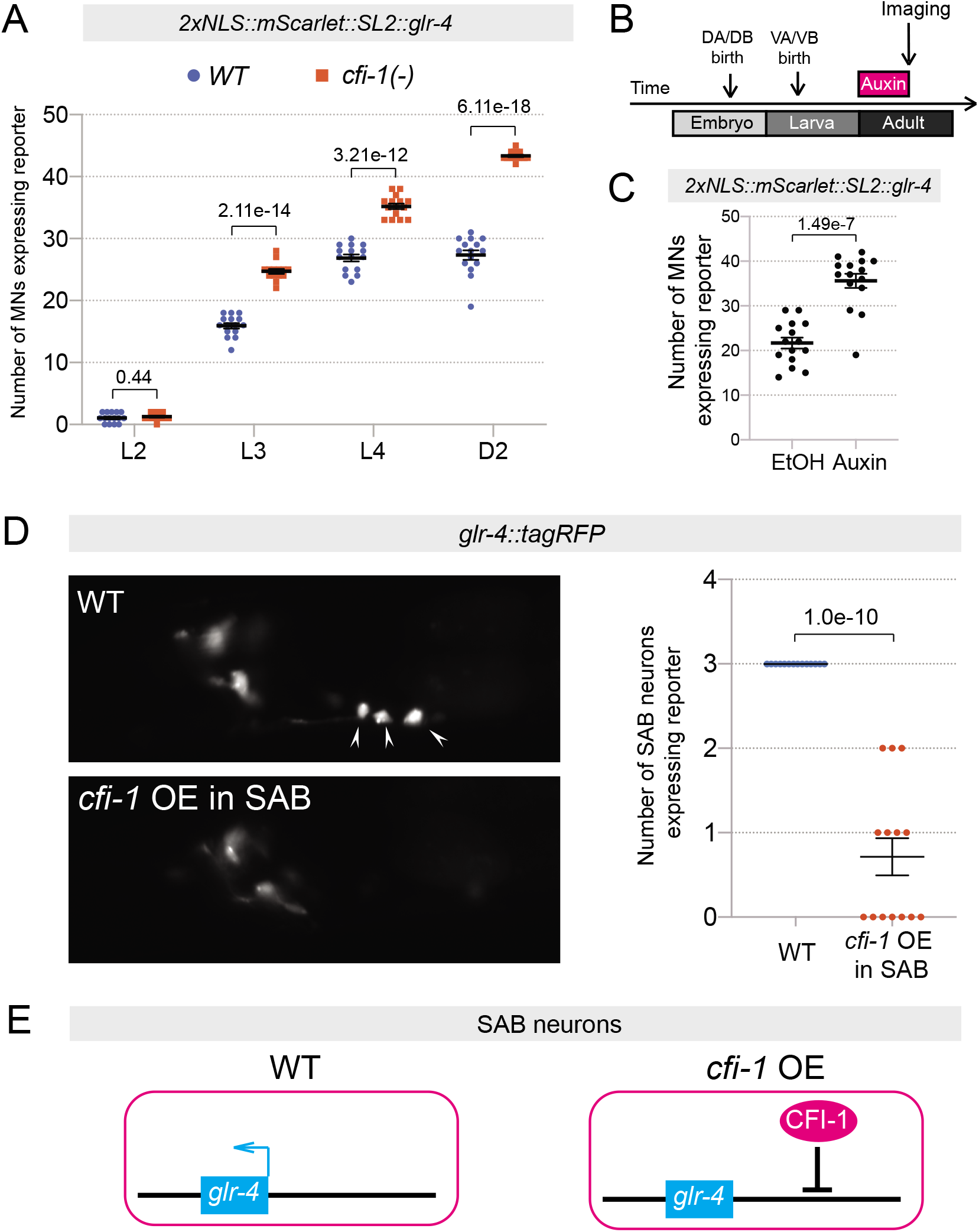
CFI-1 is sufficient to repress *glr-4* and continuously required to maintain this repression in ventral cord motor neurons. **A:** Quantification of the number of motor neurons expressing *glr-4* in WT and *cfi-1(-)* animals at four developmental stages – L2, L3, L4, and day 2 adults (D2). p-values are indicated in the graph. N ≥ 12. **B:** Diagram showing the timeline for the administration of auxin to induce degradation of CFI-1 protein in *kas16[mNG::AID::cfi-1]; otTi28[unc-11prom8+ehs-1prom7+rgef-1prom2::TIR1::mTurquoise2::unc-54 3’UTR]* animals. *otTi28* drives expression of TIR1 specifically in neurons. **C:** Quantification graph comparing expression of the endogenous *glr-4* reporter between the ethanol group (control) and auxin group on *kas16[mNG::AID::cfi-1]; otTi28[unc-11prom8+ehs-1prom7+rgef-1prom2::TIR1::mTurquoise2::unc-54 3’UTR]* animals. Ectopic expression of *2xNLS::mScarlet::SL2::glr-4* in motor neurons was detected upon CFI-1 protein knock-down. p-values are indicated in the graph. N = 15. **D:** CFI-1 is sufficient to repress *glr-4* expression. Left: representative images showing loss of *glr-4::tagRFP* expression in the SAB neurons (arrowheads) upon overexpression (OE) of *cfi-1*. Right: quantification of the number of SAB neurons expressing *glr-4::tagRFP* in wildtype and *cfi-1* overexpression animals. p-value is indicated in the graph. N ≥ 14. **E:** Model summarizing the sufficiency of CFI-1 to repress *glr-4* in SAB neurons.

To test when *cfi-1* gene activity is required for repression, we monitored endogenous *glr-4* (*mScarlet)* expression in motor neurons of WT and *cfi-1* mutant animals at larval (L2, L3, L4) and adult (day 2) stages. Compared to controls, we identified a statistically significant increase in the number of *glr-4* (*mScarlet)*-expressing motor neurons at L3, L4 and adult (day 2) stages (Fig. 4A). Next, we used a conditional *cfi-1* allele (*mNG::3xFLAG::AID::cfi-1*) that enables temporally controlled CFI-1 protein depletion upon administration of the plant hormone auxin (Li et al., 2020, Zhang et al., 2015). Depletion of CFI-1 during the first 2 days of adulthood led to a significant increase in the number of motor neurons expressing *glr-4* (*mScarlet)* (Fig. 4B-C), suggesting *cfi-1* is continuously required to maintain *glr-4/GRIK1* repression in the adult (see Discussion).

### *cfi-1/ARID3* is sufficient to repress *glr-4* expression

To test whether *cfi-1* is sufficient to repress *glr-4*, we ectopically expressed *cfi-1* in the SAB neurons. Using an SAB-specific-promoter (*unc-4*) to drive *cfi-1*, we observed a significant decrease in the number of SAB neurons expressing a *glr-4* reporter gene (Fig. 4D-E), indicating *cfi-1* is not only necessary (Fig. 3), but also sufficient to repress *glr-4* expression.

### CFI-1 binding sites proximal to *glr-4* are necessary for repression in motor neurons

CFI-1 binds to both proximal and distal *cis*-regulatory elements upstream of *glr-4* (Fig. 5A). To precisely identify the elements through which CFI-1 mediates repression, we conducted *cis-*regulatory analysis in the context of transgenic reporter animals. When *tagRFP* was driven by distal regulatory elements (2.23kb or 938bp), we did not observe differences in the number of *tagRFP* expressing motor neurons between WT and *cfi-1* mutants (Fig. 5B). However, we found an increase in the number of *tagRFP*-expressing motor neurons in *cfi-1* mutants when *tagRFP* was driven by a 3.7kb element (Fig. 5B), suggesting this element contains sequences necessary for CFI-1 repression. A translational reporter (GLR-4::GFP) driven by a 4.9kb element yielded similar results (Fig. 5B). Interestingly, a shorter *tagRFP* reporter (3.14kb) that specifically lacks the most proximal CFI-1 binding peak did not show an increase in the number of *tagRFP* expressing motor neurons in *cfi-1* mutants (Fig. 5B), suggesting proximal CFI-1 binding to the *glr-4* locus is needed for repression.

**Figure 5.**
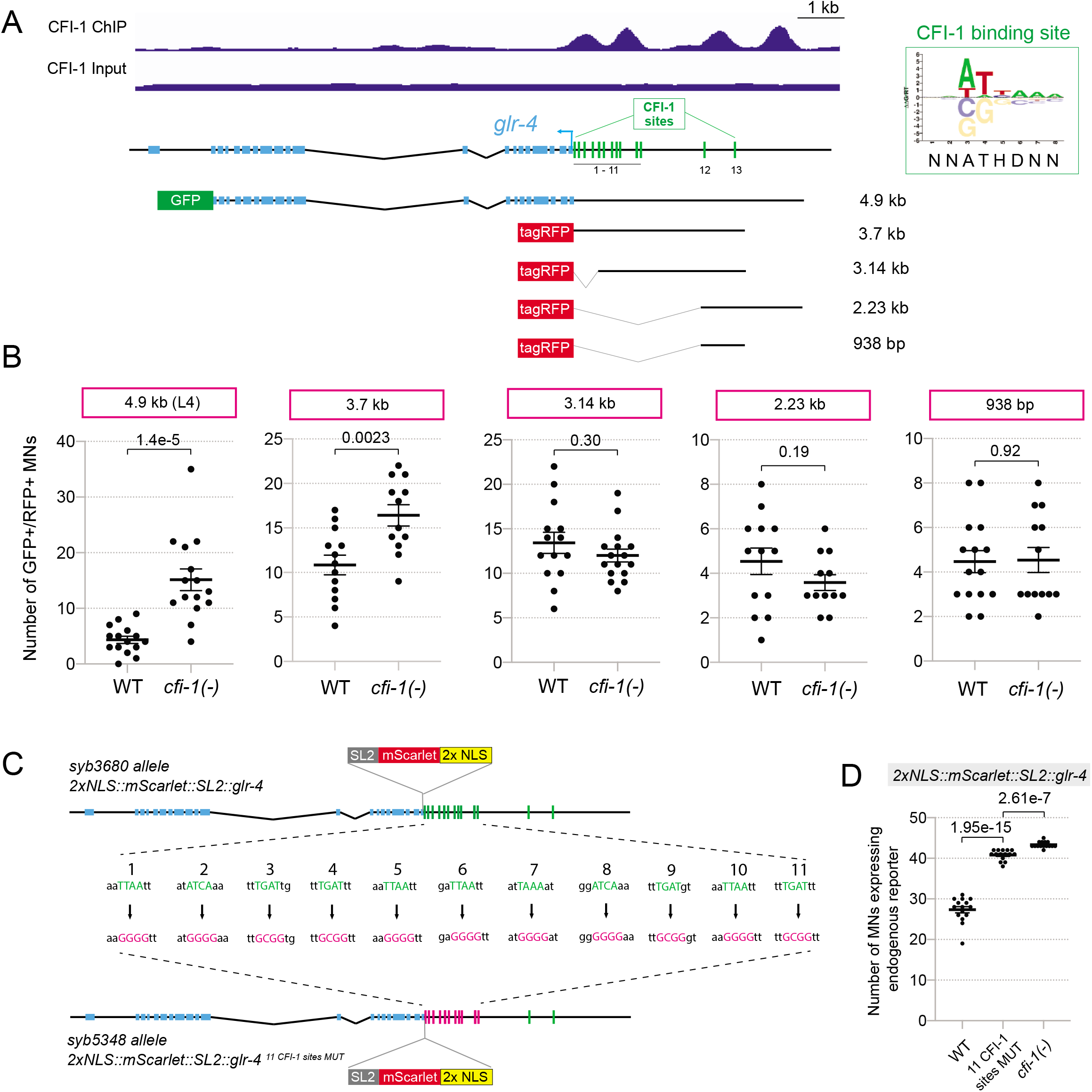
CFI-1 directly represses *glr-4* by binding to its promoter via a conserved binding motif. **A:** CFI-1 ChIP-seq tracks at the *glr-4* locus. (Bottom) Schematics of a series of *glr-4* transgenic reporter lines which contain different parts of the *cis-*regulatory regions upstream of the gene. CFI-1 recognizes a highly conserved binding motif shared by the ARID family (right). Using bioinformatic analysis, 13 of these binding motifs were identified in the *cis*-regulatory region upstream of *glr-4*, which overlap with the CFI-1 binding peaks. **B:** Quantification of the number of motor neurons expressing the reporters of *glr-4* shown in panel A in WT and *cfi-1(-)* animals in day 2 adults. Only the reporters that contain the proximal *glr-4* promoter regions show ectopic expression in *cfi-1(-)* mutants. p-values are indicated in the graph. All reporters were analyzed in the adult (D2), except the 4.9kb reporter which was analyzed at the L4 stage. N ≥ 13. **C:** Schematic depicting the design of the *2xNLS::mScarlet::SL2::glr-4^11CFI-1^ ^sites^ ^MUT^* allele. Point mutations were introduced to the 11 CFI-1 binding motifs that fall in the proximal CFI-1 binding peaks at the *glr-4* promoter. **D:** Quantification of the number of motor neurons expressing WT *2xNLS::mScarlet::SL2::glr-4* reporter and the *2xNLS::mScarlet::SL2::glr-4^11CFI-1^ ^sites^ ^MUT^*allele at day 2 adult stage. Ectopic expression of the reporter was observed with the point mutation allele, which shows a total number of *glr-4-*expressing motor neurons comparable to what was observed in *cfi-1(-)* null mutants.

We next sought to determine whether proximal CFI-1 binding sites are required for *glr-4* repression. The CFI-1 site (NNATHDNN) has been previously determined *in vitro* through protein binding microarrays (Fig. 5A) (Weirauch et al., 2014). Within the most proximal region of *glr-4*, we identified eleven predicted CFI-1 binding sites (Fig. 5A). To test their functionality, we introduced nucleotide substitutions to all eleven sites in the context of the endogenous *glr-4* (*mScarlet)* reporter through CRISPR/Cas9 genome editing (Fig. 5C). This manipulation nearly phenocopied the *cfi-1* null mutant phenotype, as it led to a dramatic increase in the number of *mScarlet*-expressing motor neurons (Fig. 5C-D). We conclude that CFI-1 binding sites located in the proximal region of *glr-4* are necessary for its repression in motor neurons.

### The transcription factor UNC-3 (Collier/Ebf) and two Hox proteins (LIN-39, MAB-5) activate basal levels of *glr-4/GRIK1* expression in motor neurons

Because *glr-4* is expressed at basal levels in cholinergic motor neurons of WT animals (Fig. 3), we reasoned this occurs due to CFI-1 antagonizing the function of *glr-4* activators in these neurons. The transcription factor UNC-3 (Collier/Ebf) and the Hox proteins LIN-39 (Scr/Dfd/Hox4-5) and MAB-5 (Antp/Hox6-8) are known to act as transcriptional activators in cholinergic motor neurons (Feng et al., 2020, Kerk et al., 2017) (Fig.6A), leading us to hypothesize that they can also activate *glr-4* expression. Indeed, we found that expression of the endogenous *glr-4* (*mScarlet*) reporter is reduced in *unc-3* loss-of-function mutant animals (Fig. 6B). Importantly, the decrease of *glr-4* expression was significantly exacerbated in *unc-3; lin-39; mab-5* triple mutants compared to *unc-3* single mutants, indicating that these three factors cooperate to activate *glr-4* in cholinergic motor neurons (Fig. 6B). Interestingly, the Hox requirement is only revealed in the absence of *unc-3* gene activity, as *glr-4* expression appears normal in *lin-39; mab-5* double mutants.

**Figure 6.**
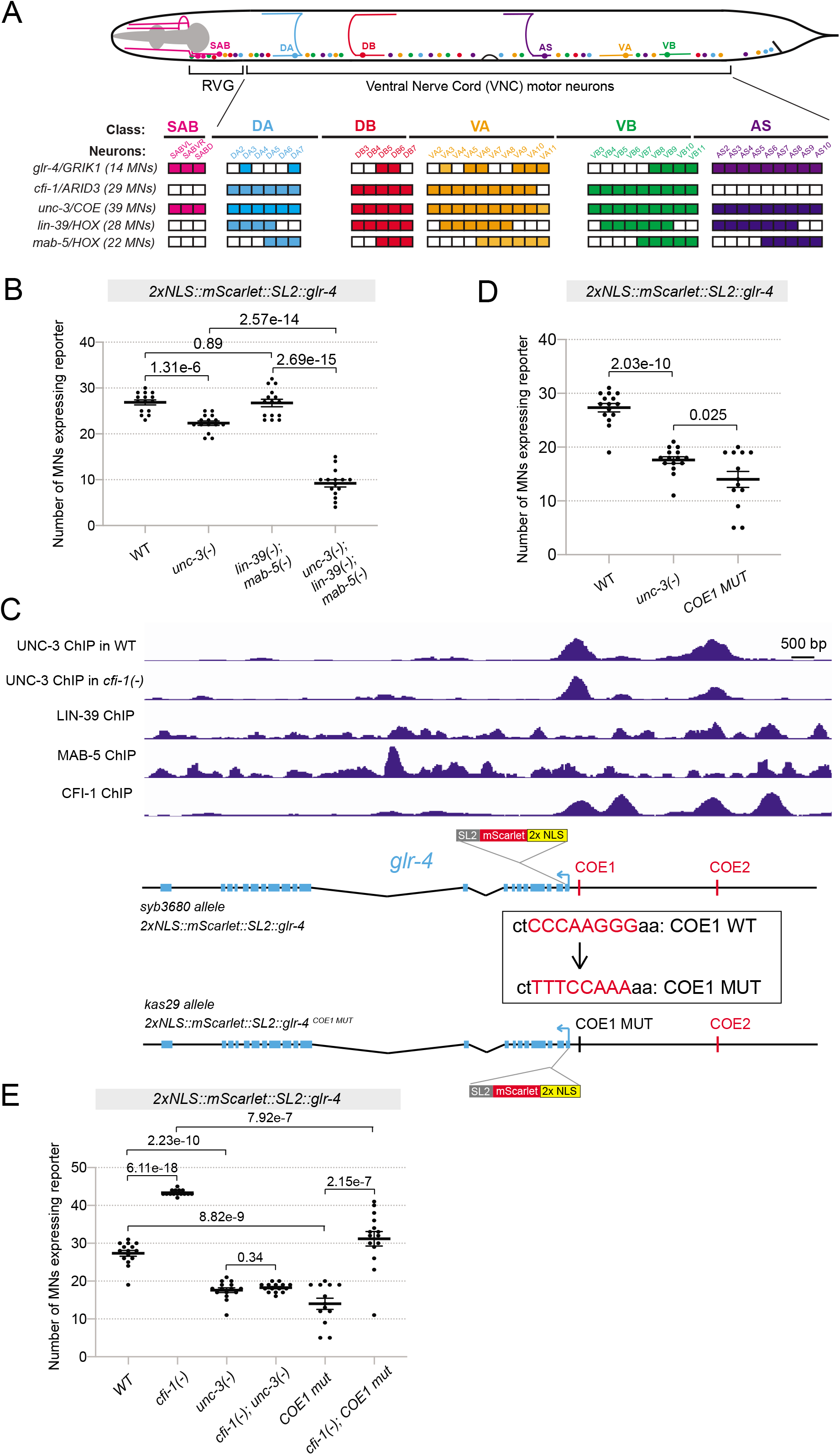
CFI-1 represses *glr-4* by counteracting activation from the cholinergic terminal selector UNC-3 and Hox proteins LIN-39 and MAB-5. **A:** Summary of the endogenous expression patterns of *cfi-1*/Arid3, *glr-4*/GRIK1, *unc-3*/COE, *lin-39*/HOX, and *mab-5*/HOX in the five cholinergic motor neuron subtypes and in the SAB neurons of the retrovesicular ganglion (RVG). Filled boxes represent positive expression, while empty boxes indicate no detectable expression. **B:** Graph summarizing quantification results of the expression of the 2xNLS::mScarlet::SL2::glr-4 reporter in WT animals, *cfi-1(-)* mutants, *lin-39(-); mab-5(-)* double mutants, and *unc-3(-); lin-39(-); mab-5(-)* triple mutants at the L4 stage. p-values are indicated in the graph. N = 15. **C:** (Doitsidou et al.) ChIP-seq binding peaks for UNC-3 (in WT animals and *cfi-1(-)* mutants), CFI-1, LIN-39, and MAB-5 at the *glr-4* locus. (Bottom) Schematic showing the details in the design of *2xNLS::mScarlet::SL2::glr-4^COE1^ ^MUT^*. Point mutations were introduced to the proximal COE motif (UNC-3 binding site) to the *glr-4* gene. **D:** Graph summarizing expression of the 2xNLS::mScarlet::SL2::glr-4 reporter in WT animals and *unc-3(-)* mutants, and the expression of the 2xNLS::mScarlet::SL2::glr-4^COE1^ ^MUT^ allele. Quantification was conducted at the day 2 adult stage. P-values are indicated in the graph. N ≥ 13. **E:** Graph summarizing expression of the *2xNLS::mScarlet::SL2::glr-4* reporter in WT animals, *cfi-1(-)* mutants, *unc-3(-)* mutants, and *cfi-1(-); unc-3(-)* double mutants and the expression of the *2xNLS::mScarlet::SL2::glr-4^COE1^ ^MUT^* allele in WT animals and *cfi-1(-)* mutants in day 2 adults. p-values are indicated in the graph. N ≥ 13.

### The most proximal UNC-3 biding site is necessary for *glr-4* expression in motor neurons

Interrogation of available ChIP-Seq data for UNC-3, LIN-39 and MAB-5 showed overlapping binding upstream of *glr-4* (Fig. 6C), suggesting a direct mode of activation by these factors. To functionally test this notion, we focused on UNC-3 because its binding site (termed COE motif) is well-defined in the *C. elegans* genome (Kratsios et al., 2012, Li et al., 2020). Through a bioinformatic search (Materials and Methods), we found two COE motifs, one proximal and one distal to the *glr-4* locus (Fig. 6C). Using CRISPR/Cas9 genome editing, we introduced nucleotide substitutions to the proximal COE motif (COE1) in the context of the endogenous *glr-4* (*mScarlet*) reporter(Fig. 6C). Animals carrying this *glr-4* (*mScarlet*) ^COE1^ ^MUT^ reporter allele showed a significant reduction in the number of *mScarlet*-expressing motor neurons, reminiscent of the effect seen in *unc-3 (-)* null mutants (Fig. 6D). These data indicate that, in WT animals, the most proximal COE motif is necessary for basal *glr-4* expression in motor neuros.

### CFI-1 antagonizes the ability of UNC-3 to activate *glr-4* expression in motor neurons

Because UNC-3 activates basal *glr-4* expression in motor neurons of WT animals, we wondered whether it also controls the increased levels of *glr-4* expression observed in *cfi-1* mutants. We found this to be the case through double mutant analysis. The number of *mScarlet*-expressing cells in *cfi-1; unc-3* mutants is dramatically decreased compared to *cfi-1* single mutants (Fig. 6E). These data indicate that CFI-1 antagonizes the ability of UNC-3 to activate *glr-4* (*mScarlet*) expression in motor neurons.

Because the proximal UNC-3 binding site (COE motif) is required to activate *glr-4* expression in WT motor neurons (Fig. 6D), this site may also be necessary for the increased expression of *glr-4* in motor neurons of *cfi-1* mutants. Indeed, we observed a significant reduction in the number of *mScarlet*-expressing cells in *cfi-1* mutants carrying the *glr-4* (*mScarlet*) ^COE1^ ^MUT^ reporters compared to the intact version of the *glr-4* (*mScarlet*) reporter (Fig. 6E). Lastly, we hypothesized that the extensive binding of CFI-1 (four binding peaks identified by ChIP-seq) immediately upstream of the *glr-4* locus may limit the ability of UNC-3 to access the locus, resulting in basal levels of *glr-4* expression in WT motor neurons. To test this, we performed ChIP-Seq for UNC-3 in WT and *cfi-1* null mutant animals (at the L3 stage). We found that UNC-3 binding on the *glr-4* locus remains largely unaltered upon *cfi-1* loss (Fig. 6C), suggesting UNC-3 can access the locus independently of the presence of CFI-1.

### The core ARID domain of CFI-1 is partially required for *glr-4* repression

To gain molecular insights into ARID3-mediated gene repression, we deleted portions of the CFI-1 DNA-binding domain and then conducted rescue assays to assess *glr-4* expression. ARID3 proteins are defined by the eARID domain, a ∼40 residue-long highly conserved domain immediately following the core ARID domain (Fig. 7A). Because structural studies on Dead ringer showed that eARID contacts DNA (Iwahara and Clubb, 1999, Iwahara et al., 2002, Patsialou et al., 2005), we tested whether the CFI-1 eARID domain is required for *glr-4* repression. Transgenic expression of either WT CFI-1 or CFI-1 lacking the eARID domain (ΔeARID) in motor neurons of *cfi-1* null mutant animals led to complete rescue. That is, *glr-4* expression was no longer observed in motor neurons when either WT or ΔeARID CFI-1 were provided (p = 0.37) (Fig. 7B), indicating eARID is dispensable for CFI-1-mediated gene repression. Next, we mutated the helix-turn-helix (HTH) domain within the core ARID region, as the HTH domain of Dead ringer contacts the major groove of DNA (Iwahara and Clubb, 1999, Iwahara et al., 2002, Patsialou et al., 2005). Again, transgenic expression of CFI-1 lacking the HTH domain (ΔHTH) in motor neurons of *cfi-1* mutants led to significant repression of *glr-4* expression -the effect is comparable to WT CFI-1 (p = 0.05) (Fig. 7B). However, transgenic expression of CFI-1 lacking the entire core ARID (ΔARID) domain (including the HTH domain) in motor neurons of *cfi-1* mutants led to partial rescue, i.e., ΔARID CFI-1 did not completely repress *glr-4* expression compared to WT CFI-1 (p = 0.0011) (Fig. 7B). Altogether, our analysis suggests that the eARID and HTH domains of CFI-1 are dispensable, but the core ARID domain is partially required for *glr-4* repression in motor neurons.

**Figure 7.**
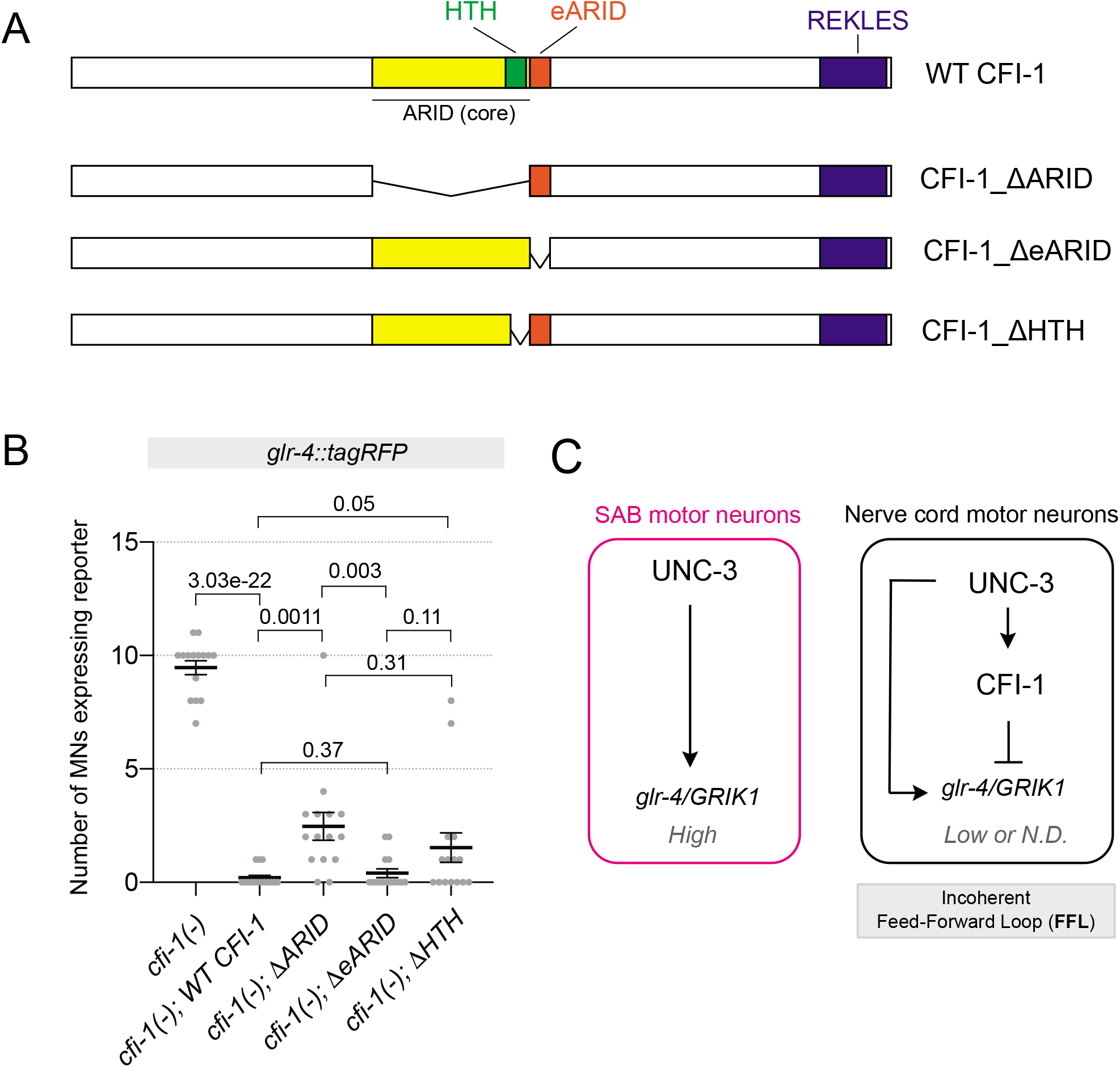
Protein motif analysis of CFI-1. **A:** Schematics of the four CFI-1 cDNA constructs tested for rescue effects including the WT cDNA, cDNA with deletion of the ARID domain (ΔARID), cDNA with deletion of the eARID domain (ΔeARID), and cDNA with deletion of the helix-turn-helix structure (ΔHTH). **B:** Quantification results of the number of motor neurons showing expression of *glr-4::tagrfp* in *cfi-1(ot786)* mutant animals expressing the rescue constructs. **C:** Schematic model summarizing our findings in SAB and nerve cord motor neurons.

## DISCUSSION

ARID3 proteins are the founding members of the ARID family, but their functions in the nervous system are poorly understood. Here, we performed ChIP-Seq for CFI-1, the sole ARID3 ortholog in *C. elegans*, and established its genome-wide binding map *in vivo*. We found that most CFI-1 target genes (77%) are expressed in post-mitotic neurons and encode proteins critical for terminal differentiation and neuronal function (e.g., neurotransmitter receptors, ion channels, neuropeptides). Further, our study offers mechanistic insights into how a single ARID3 protein controls the terminal differentiation program of distinct neuron types, uncovering cell context-depending CFI-1 functions.

### ARID3 proteins bind to proximal and distal *cis*-regulatory regions

*In vitro* assays like PBM (protein binding microarrays) and SELEX (systematic evolution of ligands by exponential enrichment) have defined the binding site of CFI-1, Dead ringer and Arid3a (Mathelier et al., 2014, Nitta et al., 2015, Weirauch et al., 2014). To assess their mechanism of action, however, unbiased assays to monitor genome-wide binding of ARID3 proteins remain necessary. ChIP-Seq in *C. elegans* for endogenously tagged CFI-1 revealed binding events predominantly located between 0 and 3kb upstream of transcription start sites (Fig. 1H). Because the size of intergenic regions for most *C. elegans* genes is less than 3kb (Dupuy et al., 2004, Nelson et al., 2004), our *in vivo* binding map strongly suggests that CFI-1 can act both at proximal (e.g., promoters) and distal (e.g., enhancers) regions to regulate gene expression. Consistently, a previous *in vitro* study found that overexpressed Arid3a in mouse embryonic stem cells also binds to both proximal and distal *cis*-regulatory elements (Rhee et al., 2014).

### The genome-wide binding map suggests a prominent role for CFI-1 in neuronal terminal differentiation

Our current knowledge of ARID3 functions in the nervous system remains rudimentary. In vertebrate nervous systems, the role of ARID3 proteins is completely unknown. In *Drosophila*, *dead ringer* has been implicated in the control of axonal pathfinding (Ditch et al., 2005, Shandala et al., 2003, Sibbons, 2004). In *C. elegans*, a handful of effector genes have been identified as CFI-1 targets in distinct neuron types (Glenwinkel et al., 2021, Kerk et al., 2017, Shaham and Bargmann, 2002, Zhang et al., 2014). Hence, a comprehensive understanding of ARID3-mediated biological processes in the nervous system is lacking.

To address this knowledge gap, we globally identified 6,396 protein-coding genes as putative direct targets of CFI-1, most of which (77%) are expressed in post-mitotic *C. elegans* neurons. GO analysis suggested a prominent role for CFI-1 in neuronal terminal differentiation, as ∼70% of its target genes encode NT receptors, transporters, ion channels, transmembrane receptors, cell adhesion molecules, etc. Moreover, 23% of CFI-1 targets encode transcription factors, chromatin factors, and proteins involved in DNA/RNA metabolism, suggesting CFI-1 can affect gene expression indirectly through these factors. Altogether, our ChIP-Seq dataset illuminates the biological processes under the control of an ARID3 protein in *C. elegans*.

### CFI-1 acts as a terminal selector in IL2 sensory neurons

Transcription factors that bind directly to the *cis*-regulatory region of terminal differentiation genes (e.g., NT biosynthesis components, NT receptors, ion channels, neuropeptides, membrane proteins) and activate their expression have been termed “terminal selectors” (Hobert, 2008, Hobert, 2016b, Hobert and Kratsios, 2019). Terminal selectors are continuously required in individual neuron types to initiate and maintain expression of terminal differentiation genes, thereby safeguarding neuronal functionality throughout life. To date, terminal selectors have been described in *C. elegans*, *Drosophila*, simple chordates and mice (Hobert and Kratsios, 2019), suggesting an evolutionarily conserved role for these critical regulators of neuronal differentiation. For most terminal selectors, however, biochemical evidence for direct binding to their target genes is currently lacking. For example, CFI-1 is a candidate terminal selector in IL2 sensory neurons because five terminal differentiation genes (*cho-1/ChT, unc-17/VAChT, gcy-19* [receptor-type guanylate cyclase], *klp-6* [kinesin-like protein], *unc-5* [netrin receptor]) fail to be properly expressed in *cfi-1* mutants (Zhang et al., 2014). Here, we provide biochemical evidence that CFI-1 binds directly to all these genes, indicating CFI-1 acts as a direct activator. Further, our analysis of the top 1,000 highly expressed genes in IL2 neurons revealed that the majority of CFI-1 binding (∼70%) occurs at terminal differentiation genes, consistent with an *in silico* prediction study for CFI-1 binding (Glenwinkel et al., 2021). Altogether, a synthesis of our findings with the aforementioned studies indicate that CFI-1 functions as a bona fide terminal selector and directly activates IL2 terminal differentiation genes. This mode of action is reminiscent of mouse Arid3a, which is known to also act as a direct activator of gene expression in cells outside the nervous system (Herrscher et al., 1995, Ratliff et al., 2014, Rhee et al., 2014).

Accumulating evidence suggests that terminal selectors act in combination with other transcription factors to determine the differentiation of individual neuron types (Glenwinkel et al., 2021, Lloret-Fernandez et al., 2018). Supporting this notion, genetic and *in silico* prediction studies found that CFI-1 collaborates with the POU homeodomain transcription factor UNC-86 to activate IL2-specific terminal differentiation genes (Glenwinkel et al., 2021, Zhang et al., 2014). Similarly, it was proposed that CFI-1 collaborates with two different homeodomain proteins, UNC-42/Prop-1 like and CEH-14/LIM, to respectively control the terminal differentiation of AVD and PVC interneurons (Berghoff et al., 2021, Glenwinkel et al., 2021), albeit the underlying mechanisms remain unclear.

### Insights into ARID3-mediated gene repression

In IL2 sensory neurons, CFI-1 exerts a dual role (Fig. 2). It functions as a direct activator of IL2-specific terminal differentiation genes and indirect repressor of ion channel-encoding genes (*pkd-2, lov-1*), which are normally expressed in CEM neurons (cells responsible for pheromone detection) (Chasnov et al., 2007). Hence, it promotes IL2 terminal differentiation and inhibits an alternative neuronal identity (CEM). However, in nerve cord motor neurons, CFI-1 functions as a direct repressor of the glutamate receptor-encoding gene *glr-4/GRIK1*. Altogether, these observations indicate that a single ARID3 protein can control the terminal differentiation program of distinct neuron types through mechanisms that depend on cell context, possibly due to CFI-1 participating in distinct neuron type-specific regulatory complexes that function either as dedicated activators or repressors.

Through an in-depth regulatory analysis of *glr-4/GRIK1*, our findings advance our understanding of how ARID3 proteins mediate gene repression in several aspects. First, *cis*-regulatory analysis combined with mutagenesis of endogenous CFI-1 binding sites strongly suggests that proximal CFI-1 binding to the *glr-4* locus is required for repression (Fig. 5). Second, CFI-1 antagonizes three conserved transcription factors (UNC-3/Ebf, LIN-39/Hox4-5, MAB-5/Hox6-8) that directly activate basal levels of *glr-4/GRIK1* expression in nerve cord motor neurons (Fig. 6). Third, ChIP-Seq for UNC-3 in WT and *cfi-1(-)* mutants did not reveal any changes in the UNC-3 binding pattern to *glr-4*, arguing against a model in which *glr-4* is repressed because the repressor (CFI-1) competes with the activator (UNC-3) for binding to the same *cis*-regulatory elements. Lastly, we found that deletion of highly conserved protein domains (eARID, HTH) predicted to bind DNA (based on Dead ringer structural studies) did not affect the ability of CFI-1 to repress *glr-4* expression (Fig. 7). This suggests that either eARID and HTH act redundantly, or other CFI-1 domains are responsible for *glr-4* repression. Supporting the latter, deletion of the entire core ARID domain (including the HTH domain) of CFI-1 led to a partial failure to repress *glr-4* in motor neurons.

One unresolved question is how exactly CFI-1 represses gene expression in *C. elegans* motor neurons. Because proximal binding is required for *glr-4* repression, it is possible that CFI-1 interferes with the function of the basal transcription complex (Baumann et al., 2010). Another possibility stems from patterning studies in the *Drosophila* embryo, where *Dead ringer* binds to Groucho, a well-characterized transcriptional corepressor (Hader et al., 2000, Valentine et al., 1998). In mouse stem cells, Arid3a directly represses pluripotency genes by recruiting histone deacetylases (HDACs)(Rhee et al., 2014). Future studies are needed to determine whether CFI-1 acts through any of these repressive mechanisms in motor neurons.

### CFI-1 continuously antagonizes the activator function of terminal selectors

The *glr-4* gene receives positive regulatory input from three conserved transcription factors (UNC-3/Ebf, LIN-39/Hox4-5, MAB-5/Hox6-8) and negative input from CFI-1. UNC-3 and the Hox proteins LIN-39 and MAB-5 are known terminal selectors in *C. elegans* nerve cord motor neurons (Feng et al., 2020, Kratsios et al., 2017, Kratsios et al., 2012). As such, they are continuously required from embryonic to adult stages to activate expression of multiple terminal differentiation genes (Feng et al., 2020, Li et al., 2020). On the other hand, our constitutive (null alleles) and conditional (temporally controlled protein depletion) approaches also revealed a continuous requirement for CFI-1 in motor neurons (Fig. 4). This argues against a transient role where, for example, CFI-1 establishes a repressive chromatin environment during early development and its activity becomes unnecessary at later stages of life. Instead, we favor a model where CFI-1 is required continuously to prevent high levels of *glr-4* expression driven by terminal selectors (UNC-3, Hox) (Fig. 7C). Such model is also supported by the continuous requirement of two other repressor proteins (BNC-1/Bnc1, MAB-9/Tbx20) in *C. elegans* motor neurons (Kerk et al., 2017). Antagonism between repressor proteins and terminal selectors has also been reported in *C. elegans* touch receptor neurons with EGL-44/Tead3 and EGL-46/Insm2 repressing terminal selector target genes (Zheng et al., 2018a). Further, studies in mice indicated that the activity of two terminal selectors, Nurr1 and Crx, which respectively control dopamine neuron and photoreceptor identities, is counteracted by repressor proteins (Otx2, Nr2e3)(Di Salvio et al., 2010, Peng et al., 2005). Additional work is needed, however, to determine whether all these repressor proteins (e.g., EGL-44, Otx2) act directly and are continuously required.

### Evolutionary implications

The terminal selector UNC-3 is required to maintain *cfi-1* expression in nerve cord motor neurons (Li et al., 2020). Hence, the repressor protein (CFI-1) and *glr-4* are both targets of UNC-3, thereby generating an incoherent feedforward loop (FFL) (Fig. 7C). In SAB motor neurons, however, CFI-1 is not expressed, and UNC-3 is able to drive high levels of *glr-4* expression (Fig. 7C)(Kratsios et al., 2015). From an evolutionary perspective, incoherent FFLs have been proposed to diversify a ground state into various substates (Hobert, 2016a). One can envision an ancestral state where an UNC-3 ortholog is present in a relatively homogeneous population of motor neurons, but the recruitment of a repressor (CFI-1) enabled their diversification. Hence, the UNC-3 → CFI-1 –I *glr-4* incoherent FFL may help distinguish, at the molecular level, nerve cord motor neurons that control locomotion from SAB motor neurons that control head movement. In agreement with this idea, incoherent FFLs are known to diversify gustatory neurons in *C. elegans* and photoreceptor cells in *Drosophila* (Etchberger et al., 2009, Johnston, 2013).

## MATERIALS AND METHODS

### *C. elegans* strain culture

Worms were grown at 20°C or 25°C on nematode growth media (NGM) plates supplied with *E. coli* OP50 as food source (Brenner, 1974). All strains used or generated for this study are listed in **Supplementary File 4**.

### Generation of transgenic animals carrying transcriptional fusion reporters and overexpression or rescue constructs

Reporter gene fusions for *cis*-regulatory analyses of *glr-4* were made with PCR fusion(Hobert, 2002). Genomic regions were amplified and fused to the coding sequence of *tagrfp* followed by the *unc-54* 3’ UTR. PCR fusion DNA fragments were injected into young adult *pha-1(e2123)* hermaphrodites at 50 ng/µl together with *pha-1* (pBX plasmid) as co-injection marker (50 ng/µl). To generate animals with *cfi-1* overexpression in the SAB neurons, the *unc-4* promoter was fused to the cDNA sequence of *cfi-1* followed by the *unc-54* 3’ UTR. The fluorescent co-injection marker *myo-2::gfp* was used (2 ng/ul) and the PCR fusion DNA fragments were injected into young adult animals carrying the *glr-4::tagrfp* reporter at 50 ng/ul. To generate transgenic animals carrying different versions of the *cfi-1* cDNA rescue constructs (WT, ΔARID, ΔeARID, ΔHTH), the *cfi-1* enhancer driving expression in motor neurons was fused to the corresponding version of *cfi-1* cDNA followed by the *unc-54* 3’ UTR. The fluorescent co-injection marker *myo-2::gfp* was used (2 ng/ul) and the PCR fusion DNA fragments were injected into young adults of *cfi-1(-)* mutants carrying the *glr-4::tagrfp* reporter at 50 ng/ul.

### Targeted genome editing

The endogenous *glr-4* reporter allele *syb3680 [2xNLS::mScarlet::glr-4]* was generated by SunyBiotech via CRISPR/Cas9 genome editing by inserting the *2xNLS::mScarlet* cassette immediately after the ATG of *glr-4*. Moreover, the endogenous *glr-4* reporter allele *syb5348* [*2xNLS::mScarlet::SL2::glr-4^11^ ^CFI-1^ ^sites^ ^MUT^*] that carries nucleotide substitutions in eleven CFI-1 binding sites was also generated by SunyBiotech. The endogenous *glr-4* reporter allele *kas29 [2xNLS::mScarlet::SL2::glr-4^COE1^ ^MUT^]* that carries nucleotide substitutions in a single UNC-3 binding site (COE1 motif) was generated in the Kratsios lab by using homology dependent repair and inserting a synthesized DNA fragment that carries the desired mutations.

### Microscopy

Imaging slides were prepared by anesthetizing worms with sodium azide (NaN_3_, 100 mM) and mounting them on a 4% agarose pad on glass slides. Images were taken with an automated fluorescence microscope (Zeiss, Axio Imager Z2). Images containing several z stacks (0.50 µm intervals between stacks) were taken with Zeiss Axiocam 503 mono using the ZEN software (Version 2.3.69.1000, Blue edition). Representative images are shown following max-projection of 2-5 µm Z-stacks using the maximum intensity projection type. Image reconstruction was performed with Image J (Schindelin et al., 2012).

### Motor neuron subtype identification

Motor neuron subtypes were identified based on combinations of the following factors: [1] co-localization with or exclusion from additional reporter transgene with known expression patterns; [2] Invariant position of neuronal cell bodies along the ventral nerve cord, [3] Birth order of specific motor neuron subtypes (e.g., during embryonic or post-embryonic stages); [4] Total cell numbers in each motor neuron subtype.

### Bioinformatic prediction of binding motifs

Information of the CFI-1 binding motif is curated in the Catalog of Inferred Sequence Binding Preferences database (http://cisbp.ccbr.utoronto.ca). To predict and identify CFI-1 binding motifs in the *glr-4* promoter, we utilized tools provided by MEME (Multiple Expectation maximization for Motif Elicitation) bioinformatics suite (http://meme-suite.org/), and performed FIMO (Find Individual Motif Occurrences) motif scanning analysis.

### Chromatin Immunoprecipitation (ChIP)

ChIP assay was performed as previously described, with the following modifications (Yu et al. 2017; Zhong et al. 2010). Synchronized L1 *cfi-1(syb1778[3xFLAG::cfi-1])* worms and N2 worms were cultured on 10 cm plates seeded with OP50 at 20°C overnight. Late L2 worms were cross-linked and resuspended in FA buffer supplemented with protease inhibitors (150 mM NaCl, 10 µl 0.1 M PMSF, 100 µl 10% SDS, 500 µl 20% N-Lauroyl sarsosine sodium, 2 tablets of cOmplete ULTRA Protease Inhibitor Cocktail [Roche Cat.# 05892970001] in 10ml FA buffer). For each IP experiment, 200 µl worm pellet was collected. The sample was then sonicated using a Covaris S220 with the following settings: 200 W Peak Incident Power, 20% Duty Factor, 200 Cycles per Burst for 1 min. Samples were transferred to centrifuge tubes and spun at the highest speed for 15 min. The supernatant was transferred to a new tube, and 5% of the material was saved as input and stored at −20°C. The remainder was incubated with FLAG antibody at 4°C overnight. Wild-type (N2) worms do not carry the 3xFLAG tag and serve as negative control. The *cfi-1(syb1778[3xFLAG::cfi-1])* CRIPSR generated allele was used in order to immunoprecipitate the endogenous CFI-1 protein. On the next day, 20 µl Dynabeads Protein G (1004D) was added to the immunocomplex which was then incubated for 2 hr at 4°C. The beads were then washed at 4°C twice with 150 mM NaCl FA buffer (5 min each), once with 1M NaCl FA buffer (5 min). The beads were transferred to a new centrifuge tube and washed twice with 500 mM NaCl FA buffer (10 min each), once with TEL buffer (0.25 M LiCl, 1% NP-40, 1% sodium deoxycholate, 1mM EDTA, 10 mM Tris-HCl, pH 8.0) for 10 min, twice with TE buffer (5 min each). The immunocomplex was then eluted in 200 µl elution buffer (1% SDS in TE with 250 mM NaCl) by incubating at 65°C for 20 min. The saved input samples were thawed and treated with the ChIP samples as follows. One (1) µl of 20 mg/ml proteinase K was added to each sample and the samples were incubated at 55°C for 2 hours then 65°C overnight (12-20 hours) to reverse cross-link. The immonuprecipitated DNA was purified with Ampure XP beads (A63881) according to manufacturer’s instructions.

### ChIP-seq data analysis

Unique reads were mapped to the *C. elegans* genome (ce10) with bowtie2 (Langmead and Salzberg 2012). Peak calling was then performed with MACS2 (minimum q-value cutoff for peak detection: 0.005) (Zhang et al. 2008). For visualization purposes, the sequencing depth was normalized to 1x genome coverage using bamCoverage provided by deepTools (Ramírez et al. 2016) and peak signals were shown in Integrated Genome Viewer (Siponen et al.). Heatmap of peak coverage in regard to CFI-1 enrichment center was generated with NGSplot (Shen et al. 2014). The average profile of peaks binding to TSS region was generated with ChIPseeker (Yu et al. 2015).

### Enrichment of CFI-1 targets in IL2 expressed genes

The top 1,000 highest expressed genes in IL2 (transcripts per million, tpm) were mined from available single-cell RNA-sequencing data (CenGEN). This dataset was computationally compared to a dataset of CFI-1 ChIP-Seq targets using the ‘semi_join’ function in R (package Dplyr 1.0.7). This generated a new data frame containing genes in the scRNA-seq dataset that are also putatively bound by CFI-1. Similarly, the ‘set_diff’ function (Dplyr 1.0.7) was used to generate a new data frame containing genes that are expressed in IL2 based on scRNA-seq but which are not found in the CFI-1 ChIP-seq dataset. Gene list analysis (PANTHER 17.0) was performed on both data frames to functionally classify all genes based on protein class ontology.

### Temporally controlled protein degradation

Temporally controlled protein degradation was achieved with the auxin-inducible degradation system (Zhang et al., 2015). TIR1 expression was driven by the pan-neuronal promoter in the transgene *otTi28[unc-11prom8+ehs-1prom7+rgef-1prom2::TIR1::mTurquoise2::unc-54 3’UTR]*. To induce degradation of CFI-1 proteins, we used the allele *kas16[cfi-1::mNG::AID].* Worms at the L4 stage were grown at 20 °C on NGM plates coated with 4 nM auxin (indole-3-acetic acid [IAA] dissolved in ethanol) or ethanol (negative control) for 2 days before testing (see figure legends for exact time in specific experiments). All plates were shielded from light.

### Statistical analysis

For data quantification, graphs show values expressed as mean ± standard error mean of animals. The statistical analyses were performed using the unpaired t-test (two-tailed). Calculations were performed using the GraphPad QuickCalcs online software (http://www.graphpad.com/quickcalcs/). Differences with p<0.05 were considered significant.

### Single molecule RNA fluorescent *in situ* hybridization (sm RNA FISH)

Synchronized L1 worms were collected from the plates and washed with M9 buffer 3 times. Worms were incubated in the fixation buffer (3.7% formaldehyde in 1x PBS) for 45 minutes at room temperature. Worms were then washed twice with 1x PBS, resuspended in 70% ethanol and left at 4°C for two nights. After removing the ethanol, worms were incubated in the wash buffer (10% formamide in 2x SSC buffer) for 5 minutes and the wash buffer was removed afterwards. A *glr-4* probe was designed using the Stellaris Probe Designer website (Biosearch Technologies). The probe was mixed in hybridization buffer (0.1 g/ml dextran sulfate [Sigma D8906-50G], 1 mg/ml *Escherichia coli* tRNA [ROCHE 10109541001], 2 mM vanadyl ribonucleotide complex [New England Biolabs S1402s], 0.2 mg/ml RNase-free BSA [Ambion AM2618], 10% formamide) and added to the worms. The hybridization buffer was removed and worms were washed twice in wash buffer (DAPI was added during the second wash and incubated for 30 minutes in the dark for nuclear counterstaining). Worms were washed once in 2x SSC, incubated in GLOX buffer (0.4% glucose, 0.1 M Tris-HCl, 2x SSC) for 2 minutes for equilibration, and the resuspended in GLOX buffer with glucose oxidase and catalase added. The samples were then examined under the fluorescent microscope.

### Real-time PCR assay for *glr-4* expression level analysis

Synchronized L4 stage wildtype and *cfi-1(-)* worms were collected, and mRNA was extracted. cDNA library was prepared using the Superscript first strand cDNA synthesis kit (Invitrogen #11904-018). RT-PCR TaqMan assays for the genes *glr-4* (assay ID: Ce02435302_g1) and *pmp-3* (Ce02485188_m1) were performed, and the expression level of *glr-4* was determined in each genotype after normalizing to the expression of the housekeeping gene *pmp-3*.

### Harsh touch behavioral assay

Harsh touch was delivered with a platinum wire pick as previously described (Marques et al., 2019). The stimulus was applied from above the animals by pressing down with the edge of the pick on the tail of non-moving adult animals. Each animal was tested only once. Worms that moved forward in response to harsh touch were scored as normal response. Results are presented as fractions of animals that responded normally.

## ACKNOWLEDGEMENTS

We thank the Caenorhabditis Genetics Center (CGC), which is funded by NIH Office of Research Infrastructure Programs (P40 OD010440), for providing strains. We are grateful to Oliver Hobert and Manasa Prahlad for comments on this manuscript. This work was funded by two NIH grants to P.K (R21 NS108505, R01 NS118078).

## AUTHOR CONTRIBUTIONS

Y. L., Conceptualization, Data curation, Investigation, Visualization, Methodology, Writing— review and editing; J.J.S. Formal analysis, Validation, Investigation, Writing—review and editing; F.M., A.O., H.C.H., Formal analysis, Validation, Investigation; P. K., Conceptualization, Supervision, Investigation, Funding acquisition, Project administration, Writing— original draft, review and editing.

## DATA AVAILABILITY

Sequencing data have been deposited in GEO under accession code GSE 205628. Moreover, all data generated or analyzed for this study are included in the manuscript and supporting files.

## LEGENDS OF SUPPLEMENTARY FIGURES AND FILES

**Supplementary Figure 1.**
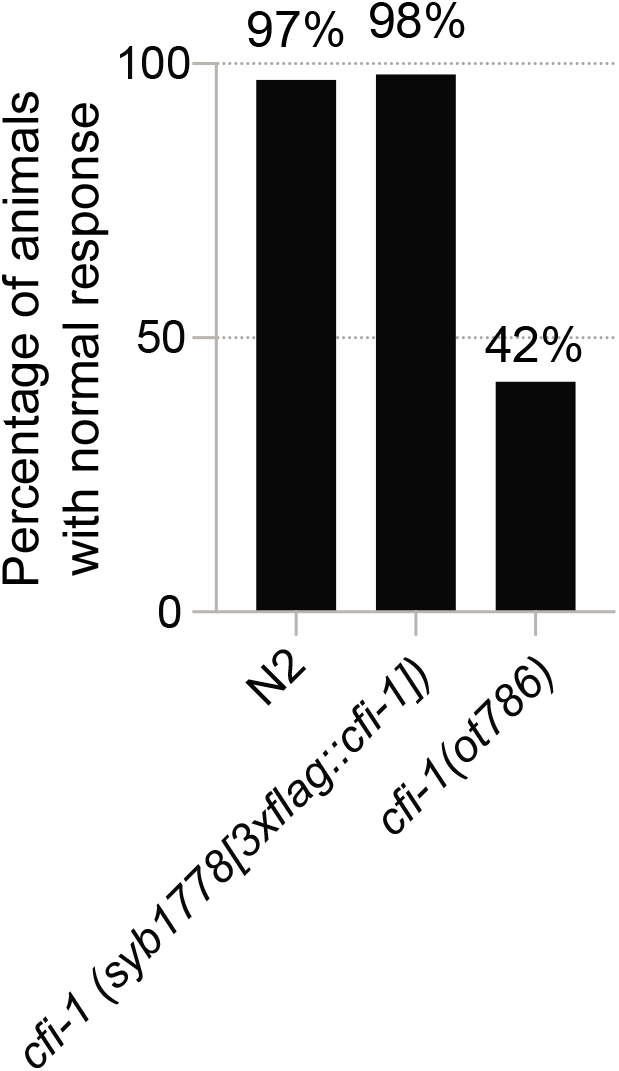
Harsh touch assay on *C. elegans* animals carrying the *3xflag::cfi-1* allele used for ChIP-Seq. Summary of the percentage of animals that show normal response to harsh touch stimulation for each genotype. Normal response is determined by forward locomotion upon harsh touch of the tail. See Materials and Methods for details. The null *ot786* allele of *cfi-1* is used as a positive control. N = 60.

**Supplementary Figure 2.**
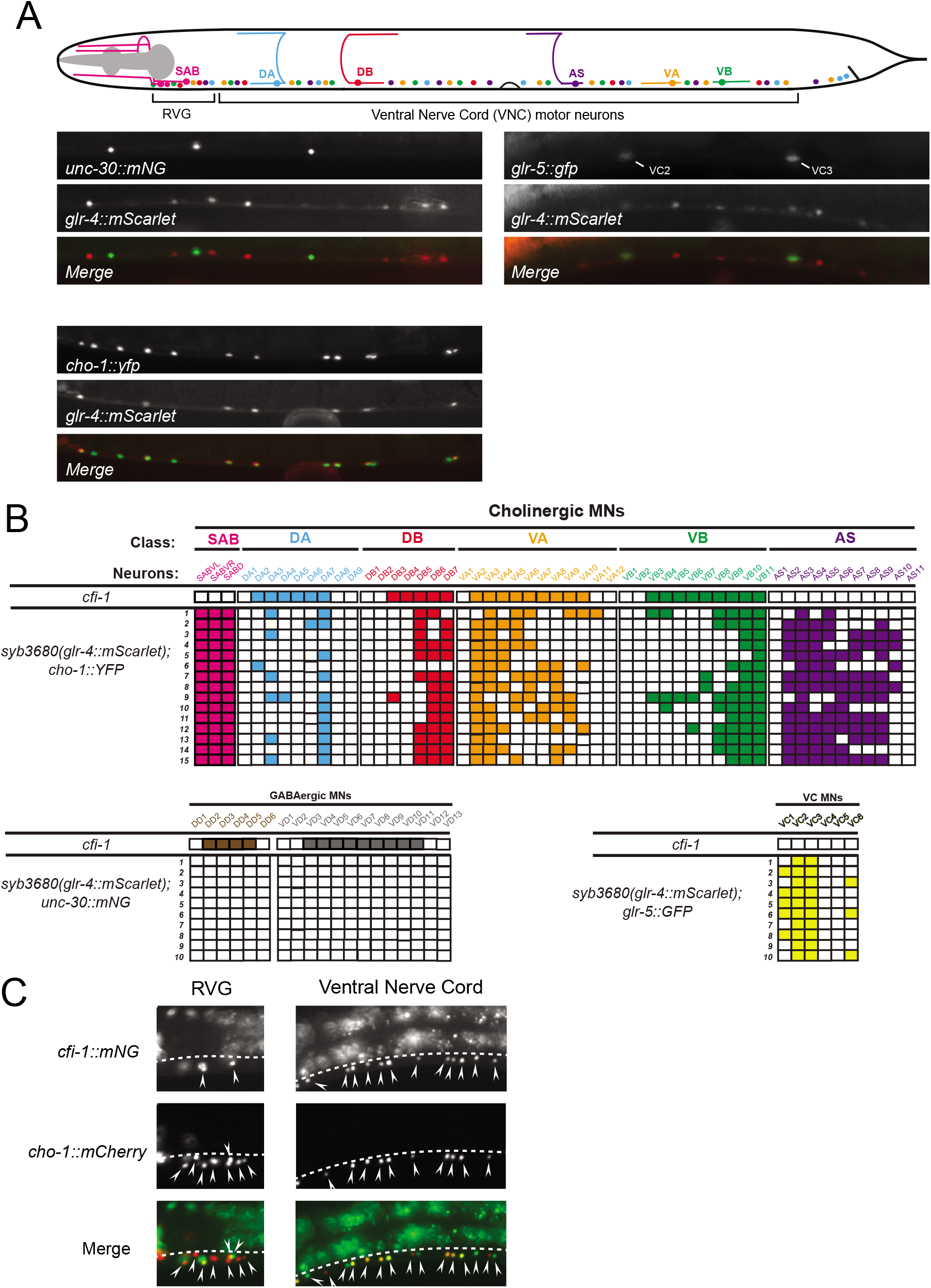
Expression of *glr-4* in nerve cord motor neurons. **A:** Representative images showing colocalization analysis of *glr-4* expression and motor neuron subtype-specific reporters *unc-30::mNG* (marker for GABAergic motor neurons), *glr-5::gfp* (marker for VC neurons), and *cho-1::yfp* (marker for cholinergic motor neurons). **B:** Summary of the expression of *glr-4* in the VNC motor neurons. Filled boxes indicate expression, while empty boxes indicate no detectable expression. Each row represents an individual worm scored at larval stage 4 (L4) for colocalization of *glr-4* and motor neuron subtype-specific reporters. **C:** Representative images showing the colocalization analysis for *cfi-1* expression in RVG and ventral cord motor neurons. The *cho-1:: mCherry* reporter is expressed in cholinergic motor neurons.

**Supplementary Figure 3.**
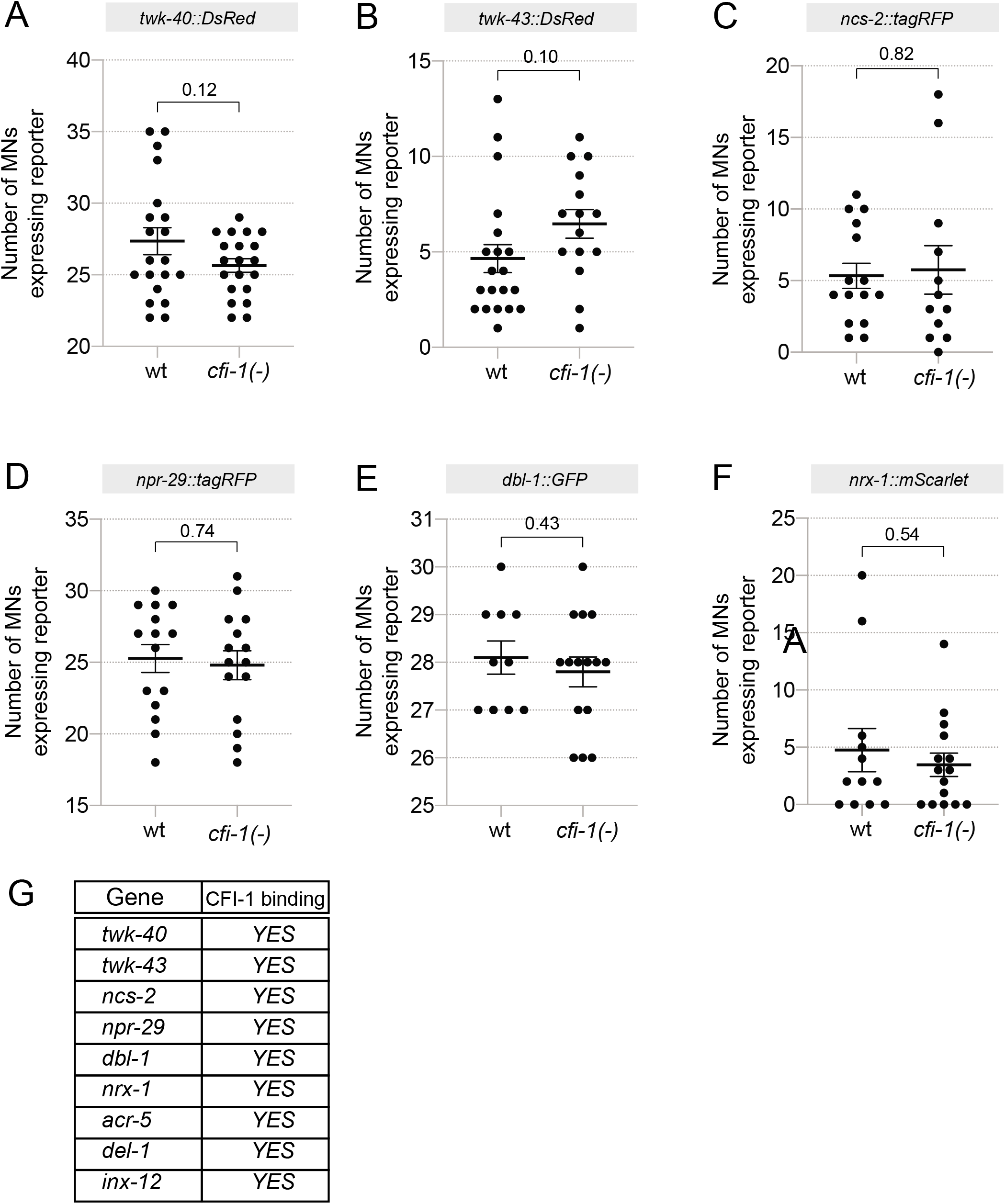
The expression of motor neuron terminal differentiation genes is not affected in *cfi-1* mutant animals. **A-F**: Quantification of the number of motor neurons expressing fluorescent reporters for terminal differentiation genes in WT and *cfi-1 (ot786)* mutant animals. Stage: L4. P-value is indicated in the graph. N ≥ 10. **G**: All genes are bound by CFI-1 based on ChIP-seq data.

**Supplementary File 1.** Genes expressed in *C. elegans* neurons and bound by CFI-1.

**Supplementary File 2.** Protein ontology analysis on genes bound by CFI-1.

**Supplementary File 3.** Protein ontology analysis on genes expressed in IL2 neurons and bound by CFI-1.

**Supplementary File 4.** List of *C. elegans* strains used or generated for this study.

